# Probing the eukaryotic microbes of ruminants with a deep-learning classifier and comprehensive protein databases

**DOI:** 10.1101/2024.07.17.603995

**Authors:** Ming Yan, Thea O. Andersen, Phillip B. Pope, Zhongtang Yu

## Abstract

Metagenomics, particularly genome-resolved metagenomics, has significantly deepened our understanding of microbes, illuminating their taxonomic and functional diversity and roles in ecology, physiology, and evolutions. However, eukaryotic populations within various microbiomes, including those in the mammalian gastrointestinal (GI) tract, remain relatively underexplored in metagenomic studies due to the lack of comprehensive reference genome databases and robust bioinformatics tools. The GI tract of ruminants, particularly the rumen, contains a high eukaryotic biomass although a relatively low diversity of ciliates and fungi, which significantly impact feed digestion, methane emissions, and rumen microbial ecology. In the present study, we developed GutEuk, a bioinformatics tool that improves upon the currently available Tiara and EukRep in accurately identifying metagenome eukaryotic sequences. GutEuk is optimized for high precision across different sequence lengths. It can also distinguish fungal and protozoal sequences, facilitating further elucidation of their unique ecological and physiological impacts. GutEuk was shown to facilitate a comprehensive analysis of protozoa and fungi within more than one thousand rumen metagenomes, revealing a greater genomic diversity among protozoa than previously documented. We further curated several ruminant eukaryotic protein databases, significantly enhancing our ability to distinguish the functional roles of ruminant fungi and protozoa from those of prokaryotes. Overall, the newly developed package GutEuk and its associated databases create new opportunities for in-depth study of GI tract eukaryotes.

## Introduction

The complex and multi-kingdom microbial ecosystem unique to the rumen supplies ruminants with up to 70% of their energy requirement through the digestion and fermentation of plant materials^1^. However, this process also results in methane emissions, which account for approximately 14% of the anthropogenic greenhouse gas emissions^2^. The rumen microbes also convert dietary nitrogen, including protein and non-protein nitrogen, into high-quality microbial protein, contributing up to 80% of the protein metabolized in the small intestine^3^. While the rumen microbiome is primarily linked to ruminant nutrition and production, the lower gastrointestinal (GI) microbiome is more associated with animal health and host physiology^4^. The intricate and interdependent multi-kingdom ecosystem comprises anaerobic bacteria, archaea, fungi, protozoa, and viruses, each occupying a unique niche. While the prokaryotic communities of the rumen and intestines have been extensively investigated through multi-omics studies, analysis of the eukaryotic communities still primarily relies on amplicon sequencing and analysis of a phylogenetic marker (metataxonomics), mainly the 18S rRNA gene for protozoa and internal transcribed spacer (ITS) for fungi, which provide limited functional information. The lack of a comprehensive genome database for rumen protozoa and fungi further limits omics analyses of these eukaryotes. Our understanding of the functional importance of the eukaryotic communities within the rumen is still primarily based on culture-based and genomic studies of a small number of species.

Unlike the human gut, where eukaryotes account for less than 1% of the gut microbiome^5^, fungi and protozoa constitute up to 20% and 50% of the rumen microbiome biomass respectively^4,6,7^. Rumen fungi are specialized fiber degraders capable of producing diverse carbohydrate-active enzymes (CAZymes), especially those hydrolyzing recalcitrant plant cell wall materials^8^ and complementing the CAZymes repertoire of rumen bacteria^8,9^. In addition, anaerobic fungi produce diverse secondary metabolites with potential antimicrobial and therapeutic properties^10^. Except *Pecoramyces sp*. F1, which was isolated from the rumen^11^, the available genomes of anaerobic fungi of ruminants were acquired exclusively from the feces, explicitly *Anaeromyces robustus, Caecomyces churrovis, Neocallimastix californiae, Neocallimastix lanati*, and *Pecoramyces ruminantium*, therefore also representing hindgut fungi. Although the rumen and the hindgut share some of their fungi, they have distinctive fungal communities^12^, and it remains unknown whether the fungal enzymatic repertoire differs between the rumen and the hindgut. While the contribution of fungi to fiber degradation in the rumen is well-recognized, the functions of rumen protozoa are intricate and multifaceted, encompassing productive roles such as fiber degradation and rumen pH stabilization, alongside counterproductive roles such as promoting methane emissions and wasteful intra-ruminal recycling of microbial protein^13^. Indeed, rumen protozoa exhibit a linear relationship with methane emissions^14^. Due to their negative associations with feed efficiency and methane emissions, a few studies have attempted to specifically inhibit the dominant species of *Entodinium*^15,16^. Alternatively, analyses of single-cell amplified genomes (SAGs) recovered from the rumen and subsequent proteomic analysis of rumen protozoa revealed an unexpectedly extensive repertoire of various CAZymes^7,17^. Recent studies showed a relationship between rumen protozoa and feed efficiency in beef cattle^18^ and between intestinal protozoa and calve intestinal health^19^. Therefore, it is imperative to transcend beyond metataxonomic analysis of rumen protozoa, characterizing the genomes and enzymatic capacity of the eukaryotic fraction in the GI tract of ruminants. Moreover, a more comprehensive protein database is required for robust, unbiased analyses of gene/protein expression profiles of protozoa in the context of important rumen functions and animal production, such as methane emissions and feed efficiency.

Single-cell genomics has proven valuable in unraveling the fundamental biological characteristics of protozoa^17^. However, it is not cost-effective for ecological studies of protozoal communities. Metagenomics has revolutionized the investigation of various microbiomes, including the GI microbiomes of ruminants, but most studies focus on bacteria and archaea while ignoring fungi and protozoa. Two machine learning-based tools (Tiara and EukRep) are available to identify eukaryotic sequences directly from metagenomic assemblies^20-22^. Studies using these tools have revealed the ecological importance of protists in aquatic environment^23-26^. However, it remains unclear whether the above tools can be used to analyze eukaryotes abundant in host-associated ecosystems like the rumen. Moreover, the diversity and functional repertoire of the ruminant GI tracts, especially the less-studied hindgut and less-represented ruminant species, remains limited. Here, we benchmarked the existing tools for classifying ruminant gut eukaryotes and created a new tool (referred to as GutEuk) to identify and classify eukaryotic microbes (fungi and protozoa) in metagenomes derived from the gut of ruminants. Using the newly developed tool, which surpasses the existing tools in both precision and recall, we reanalyzed over one thousand metagenomes and metatranscriptomes from ruminant GI tracts. This analysis included samples from foregut (mostly rumen but also including reticulum, omasum, and abomasum) and hindgut (cecum, colon, and rectum) across various ruminant species (mostly domestic species including *Bos taurus* and *Ovis aries* as well as the wild species including *Capreolus capreolus* and *Hydropotes inermis*) and led to the creation of protein databases for both ruminant foregut and hindgut eukaryotes. Using the newly established protein databases, we reanalyzed the metaproteomes from a recent study^7^ and observed a significant increase in the number of proteins identified as fungal or protozoal. GutEuk and the new protein databases open up new opportunities in analyzing and reanalyzing host-associated microbiomes for eukaryotic microbes, especially those within the rumen.

## Results

### The new GutEuk improves identification of eukaryotic microbes from metagenomes

Tiara^20^ and EukRep^21^ are tools for identifying eukaryotic sequences from metagenomic assemblies based on *k*-mer frequency. They were developed based on a limited selection of eukaryotic nuclear genomes (73 for Tiara and 70 for EukRep), encompassing those of algae and animals. The lack of representation of gut-associated eukaryotic microbes likely undermines their utility in analyzing gut-derived metagenomes. In addition, *k*-mer frequency merely represents metrics of simplified genomic contexts, while improved classification can be achieved using convolutional neural network (CNN) with original DNA sequences as inputs, as illustrated by the recent study^27^. We tested Tiara and EukRep for their ability to identify gut eukaryotic sequences using a dataset from 52 SAGs of rumen protozoa^17^ (see Methods). Not so surprisingly, Tiara and EukRep correctly classify only 30,081 and 44,395 out of 57,253 eukaryotic contigs, achieving accuracies of 52.5% and 77.5%, respectively, which are much lower than their reported performance.

Therefore, these two existing tools cannot achieve a generalizable performance for less-represented genomes of eukaryotes. We thus developed a specialized deep-learning classifier built upon diverse protozoal, fungal, and prokaryotic genomes to improve the classification of eukaryotic microbes in the rumen and gut. Leveraging thousands of prokaryotic, fungal, and protozoal genomes, we developed GutEuk, an ensembled deep-learning model combining CNN (one-hot encoded nucleotide bases as inputs) and feedforward neural network (FNN; *k*-mer frequencies as inputs) (Fig. 1a and 1b). GutEuk employs a two-stage classification with user-defined confidence levels at each stage: The first stage differentiates between prokaryotic and eukaryotic (fungal and protozoal) sequences and the second stage further distinguishes fungal from protozoal sequences.

**Fig. 1:**
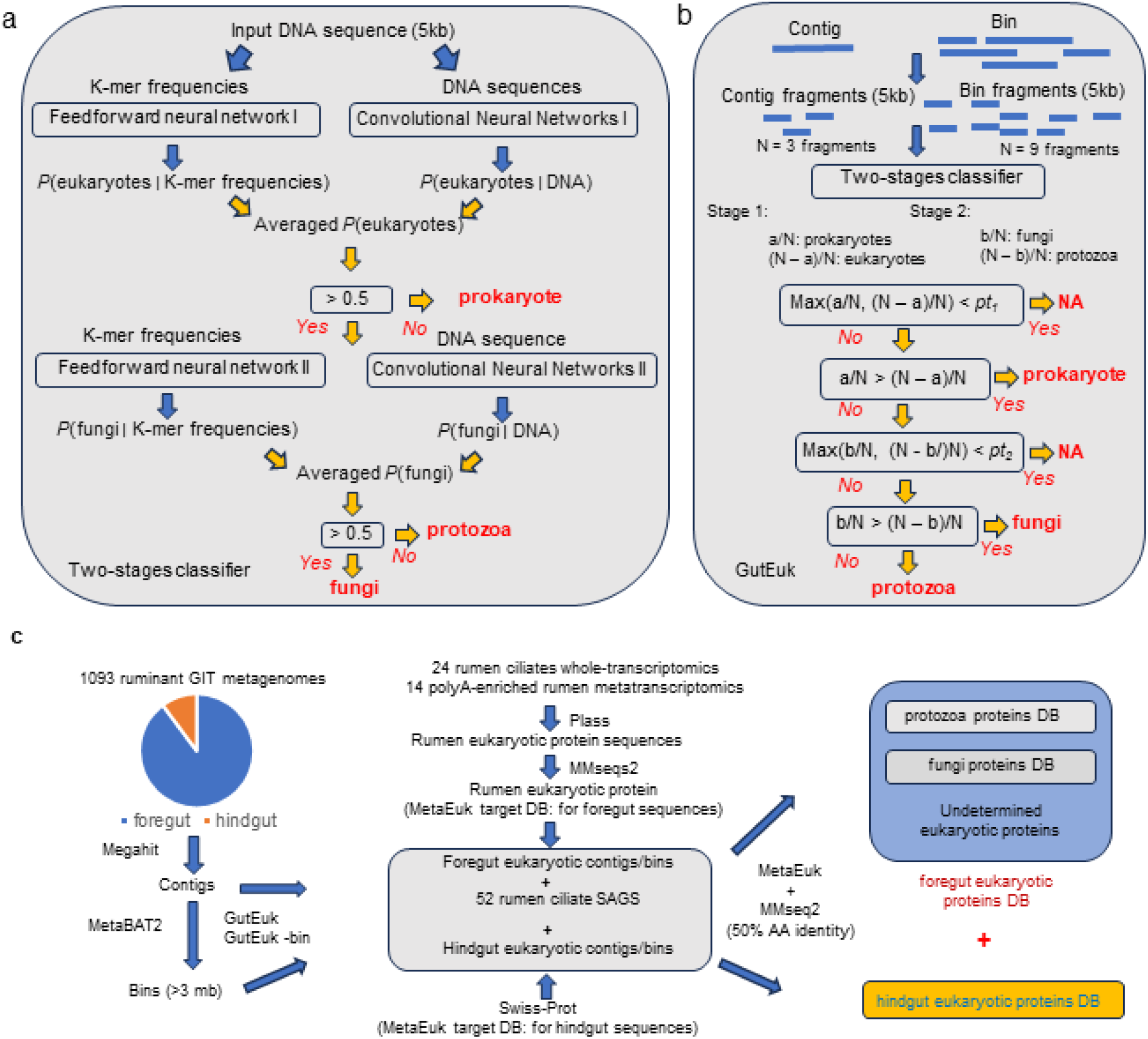
The schematic of the two-stage classification process used by GutEuk. (**a**), the pipeline to classify individual contig and bin from the metagenomes (**b)**, the workflow to establish the ruminant foregut and hindgut eukaryotic protein databases.

We benchmarked the performance of GutEuk against EukRep and Tiara using independent datasets not used for training of these tools at the contig level (see Methods). The results showed that GutEuk outperformed Tiara and EukRep consistently at different contig lengths considerably (Fig. 2a). GutEuk and Tiara exhibited comparable precision for eukaryotic contigs, while EukRep displayed a slightly lower precision, especially for contigs shorter than 20kb. However, for prokaryotic contigs, GutEuk achieved a precision of around 99% consistently, even for contigs shorter than 15kb, whereas Tiara and EukRep achieved a precision of 95% and 91%, respectively, for contigs shorter than 10kb. The performance of Tiara and EukRep was poorer for contigs between 3 and 5kb. In terms of recall rate, all three tools performed similarly with prokaryotic contigs, while for eukaryotic contigs, GutEuk achieved a recall rate of around 99% for contigs shorter than 15kb and still maintained a recall rate of around 98% for longer contigs. In comparison, EukRep and Tiara recorded a lower recall rate: 95% at best and less than 90% with contigs shorter than 10kb for EukRep, while for Tiara, only 82% at best with contigs between 5 and 10kb and lower recall rates with longer contigs.

**Fig. 2:**
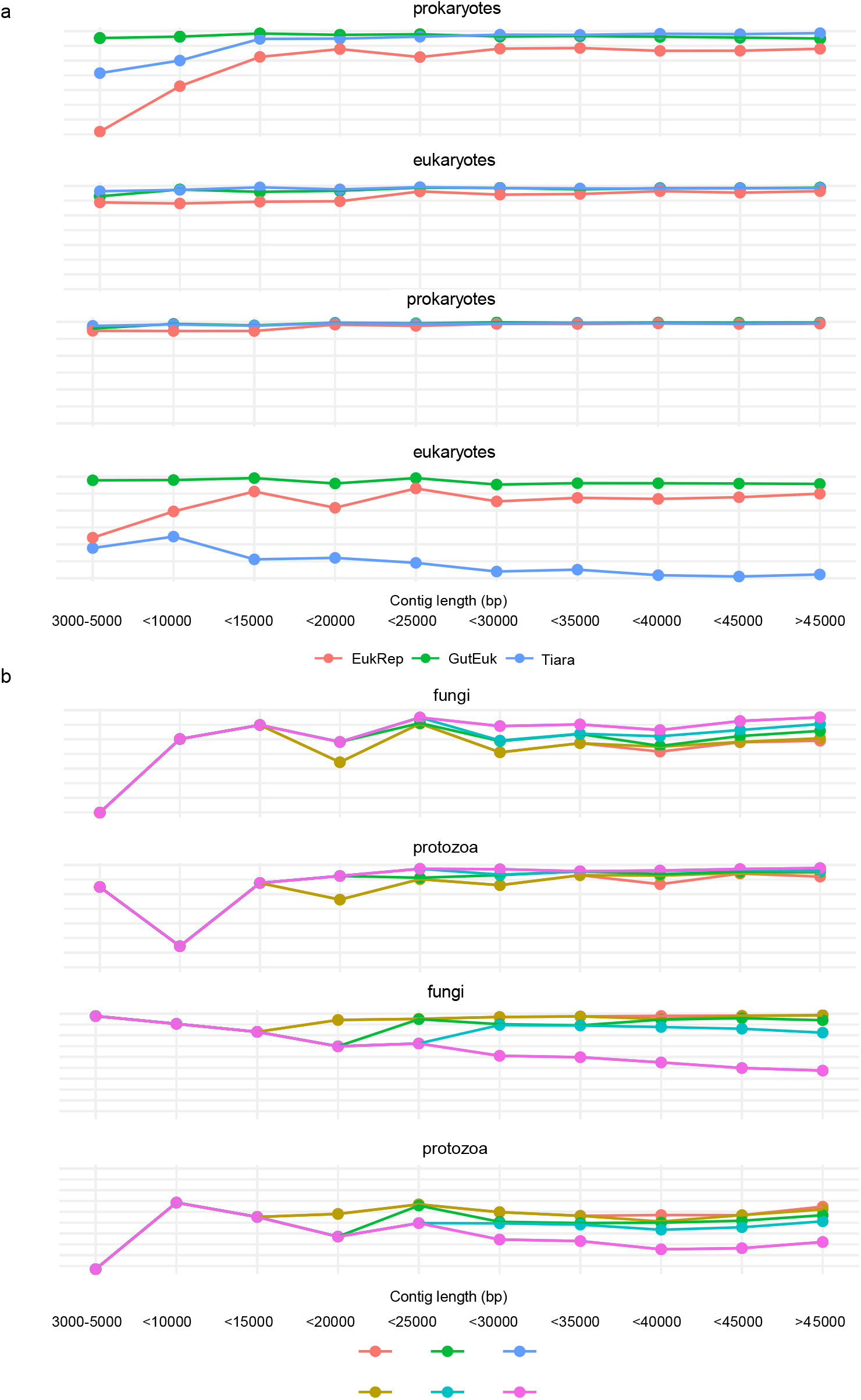
Benchmarking the performance of GutEuk, EukRep, and Tiara for contigs. **a**, The performance of GutEuk, EukRep, and Tiara in differentiating prokaryotic and eukaryotic contigs at different lengths. EukRep and Tiara were implemented at the default setting. GutEuk was implemented with the *pt*_*1*_ of 0.5. **b**, The performance of GutEuk in further differentiating fungal and protozoal contigs at different confidence levels (*pt*_*2*_) and contig lengths.

We evaluated GutEuk for its ability to further differentiate protozoal and fungal contigs (Fig. 2b). Since for contigs shorter than 15kb, the maximum number of classifiable fragments is two, the threshold used will not affect its performance (both fragments must provide the same prediction). For contigs exceeding 15kb, we observed the expected increase in precision and decrease in recall as the confidence level (*pt*_*2*_) rose. Specifically, a precision above 90% was seen even for *pt*_*2*_ of 0.5, and a 95% precision was seen for *pt*_*2*_ above 0.8. In terms of recall rate, GutEuk achieved above 70% and 85% for protozoa and fungi, respectively, when a *pt*_*2*_ of 0.8 was used. Therefore, *pt*_*2*_ can be adjusted to optimize GutEuk for precision or recall to address specific needs or questions. Nevertheless, a balance between precision and recall was seen with a *pt*_*2*_ of 0.8, and the corresponding precision of above 95% was considered conservative.

In addition to evaluating the performance in identifying microbial eukaryotic contigs, we further investigated whether GutEuk could achieve consistent results across genomes of diverse phylogenetic lineages, including those of aquatic fungi and protozoa included in the database (Fig. 3). All three tools achieved above 90% accuracy. However, the expanded training set and the more complex model used in GutEuk allowed it to achieve over 90% accuracy in classifying DNA from most eukaryotic microbes, especially the underrepresented rumen ciliate protozoa. The only eukaryotic genome whose fragments were identified with a lower than 70% accuracy (about 45%) was the genome of *Stentor coeruleus* (accession id: GCA_001970955.1), a freshwater protozoan. In contrast, Tiara and EukRep each had 14 eukaryotic genomes classified with an accuracy lower than 70%. These results show that at a *pt*_*1*_ of 0.5, GutEuk predicts eukaryotic microbial genomes/bins more accurately than Tiara and EukRep. More importantly, GutEuk can further classify eukaryotic genomes as fungal or protozoal with high accuracy, except for a few fungi and protozoa not found in the mammalian GI tract. Specifically, GutEuk identified the following fungal genomes with lower than 50% accuracy: *Smittium mucronatum* (GCA_001953115.1, an insect gut symbiont), *Vavraia culicis* (GCA_000192795.1, a microsporidian parasite), and *Enterospora canceri* (GCA_002087915.1, infecting mosquito and European shore crab). Similarly, the protozoal genomes identified with less than 50% accuracy are not found in GI tract either: *Perkinsus marinus* (GCA_000006405.1) and *Perkinsus* sp. *BL_2016* (GCA_004369235.1), two parasitic flagellate inflecting mollusks, as well as *Stentor coeruleus* (GCA_001970955.1) and *Lenisia limosa* (GCA_001655205.1), two aquatic ciliates. Given that gut eukaryotes exhibit a narrow phylogenetic range, GutEuk is expected to perform well with gut metagenomic sequences (see the next section). A higher *pt*_*2*_ threshold (e.g., 0.8) can be used if a lower false positive rate is required.

**Fig. 3:**
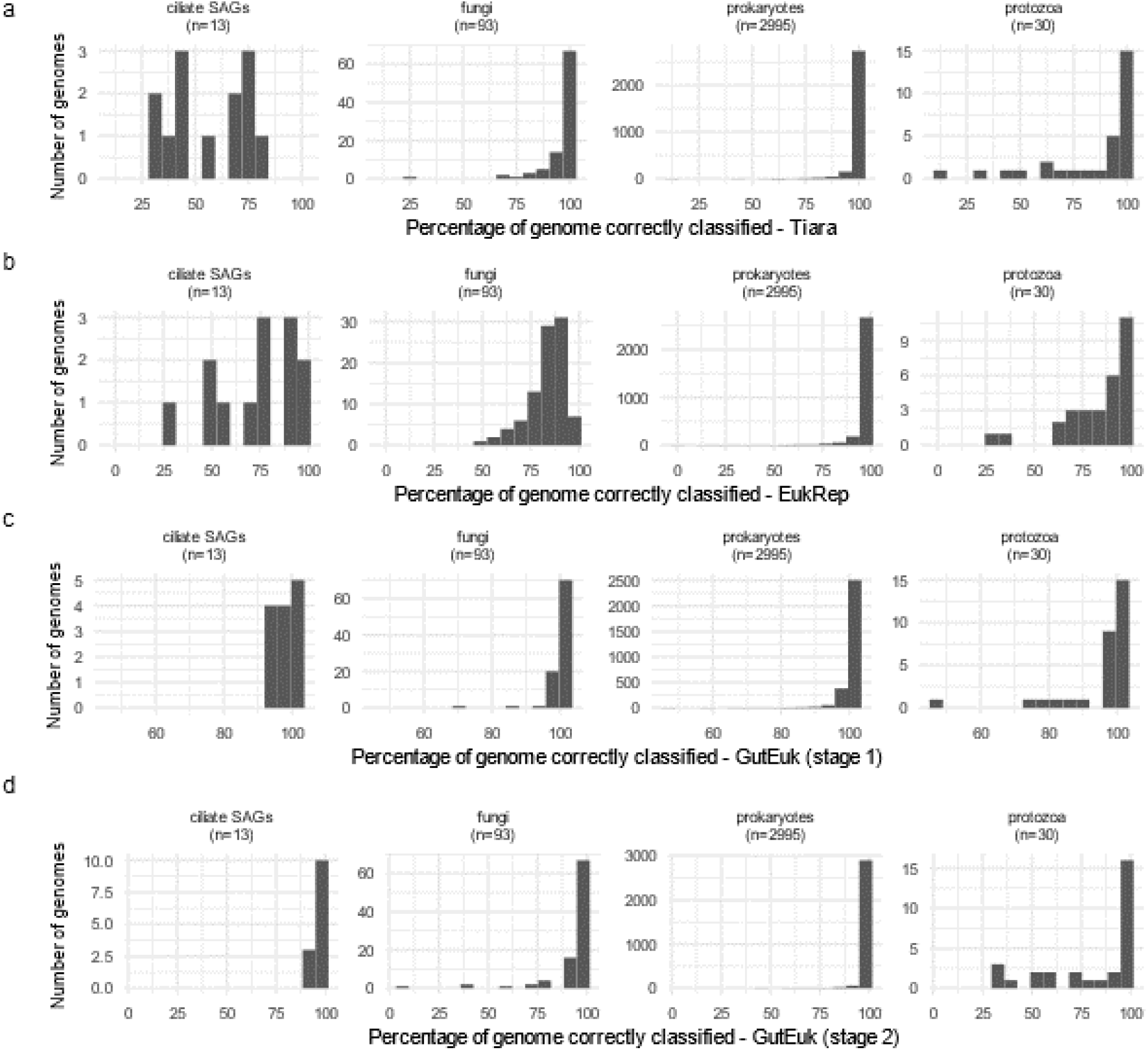
Benchmarking the performance of GutEuk, EukRep, and Tiara for bins. The percentage of genomes being correctly classified as prokaryotic or eukaryotic by Tiara (**a**), EukRep (**b**), and GutEuk (**c**). The percentage of genomes being correctly classified as prokaryotic, protozoal, or fungal by GutEuk (**d**).

### Diverse rumen protozoa are identified from underrepresented ruminant species

Until now, 53 genomes have been sequenced, including 52 SAGs^17^ and one mono-species genome^28^. These genomes represent 19 species of rumen ciliates prevalent across various ruminant species. Using GutEuk, we systematically analyzed the 15,000 bins (each greater than 3 MB) assembled from 1,093 previously sequenced rumen metagenomes (Supplementary Table 1, see Methods) for eukaryotic bins. These samples were collected from 13 different ruminant species, encompassing both domestic and wild species with varying feeding regimes. We identified hundreds of protozoal bins and one fungal bin. The scarcity of bins identified as fungal likely reflects their low abundance and low fungal DNA yield from rumen fluid samples because rumen fungi strongly adhere to digesta particles. We assessed the genome completeness of the identified eukaryotic bins based on the presence and absence of lineage-resolved single-copy marker genes. Unfortunately, most of the bins were deemed largely incomplete, consistent with findings from a recent study using EukRep for eukaryotic bin identification^25^.

Despite the majority of the eukaryotic bins being incomplete, we identified 21 protozoal bins with more than 50% completeness and less than 10% contamination. We further conducted a phylogenetic analysis on this set of protozoal bins and the rumen protozoa SAGs (Fig. 4). Examination of the phylogenetic tree, derived from concatenated 30 single-copy marker proteins, reveals that the newly discovered rumen protozoal bins cluster alongside the SAGs within the *Vestibuliferida* order. In contrast, all *Entodiniomorphida* SAGs form a distinct cluster, suggesting the potential for a greater diversity of *Vestibuliferida* species yet to be uncovered. Notably, the phylogenetic tree constructed with the 30 single-copy marker proteins exhibited the same tree topology as the original phylogenetic tree of the SAGs, which was based on 113 single-copy marker proteins. This consistency demonstrates the robustness of our phylogenetic analysis. It is also noteworthy that some of the newly identified protozoal bins were derived from metagenomes of ruminant species from which no protozoal genomes have been sequenced, including *Bison bison* and *Bos indicus*. Therefore, future protozoal research should expand to include less-studied ruminant species to facilitate a more comprehensive genomic representation of rumen eukaryotes.

**Fig. 4:**
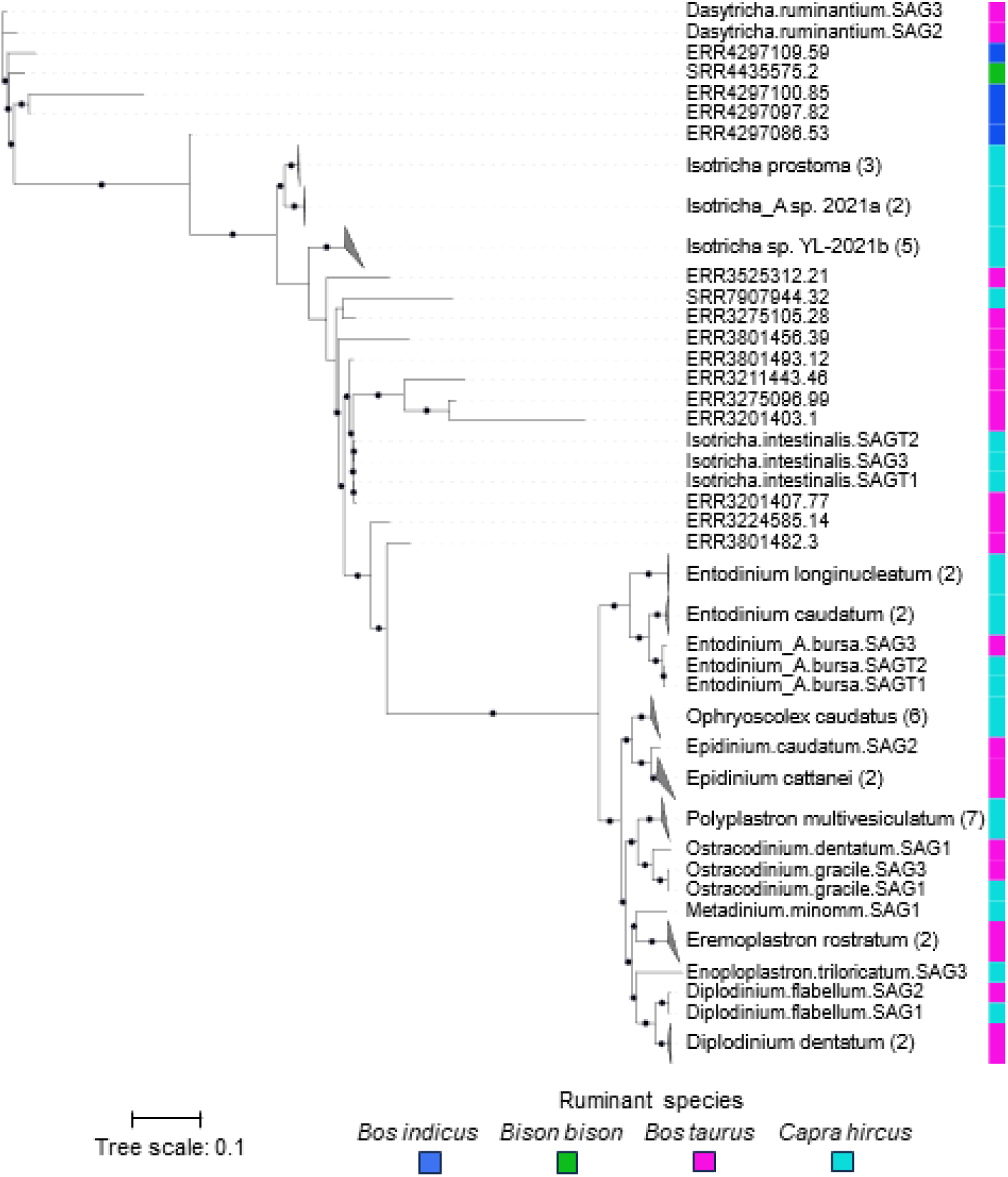
A phylogenomic tree of the identified rumen protozoal bins and 49 rumen protozoa single-cells assembled genomes. All nodes with a greater than 85% bootstrap support are indicated with dots in the branch.

### Protein databases of ruminant gut eukaryotes and the repertoire of genes encoding glycoside hydrolases and peptidases

From the eukaryotic sequences (bins and contigs) identified from the gut metagenomes of various ruminants, predominantly cattle, sheep and goats, we predicted and translated their protein sequences, subsequently constructing several protein databases (Fig. 1c). These gut eukaryotic microbial protein databases include one each for foregut protozoa (both newly identified and from the 52 SAGs), foregut fungi, and hindgut eukaryotes (Fig. 5a). The foregut protozoal database consisted of 25,498,129 proteins, which were clustered at 50% amino acid identity into 5,265,119 protein clusters, 54% of which are newly identified. Among these clusters, only 10% were annotated. The foregut fungal database is much smaller, containing 11,314 proteins, which formed 7,293 protein clusters. Around half of the fungal protein clusters could be annotated. Notably, the proportion of the annotated hindgut eukaryotic protein clusters exceeded 92%. The significantly smaller proportion of annotated protozoan proteins, compared to those of fungal and hindgut eukaryotes, highlights the need for additional genomic studies on rumen protozoa. This need is also underscored by their complex behaviors, such as predation on other rumen microbes, enzymatic breakdown of plant material, and symbiotic interactions with bacteria and archaea.

**Fig. 5:**
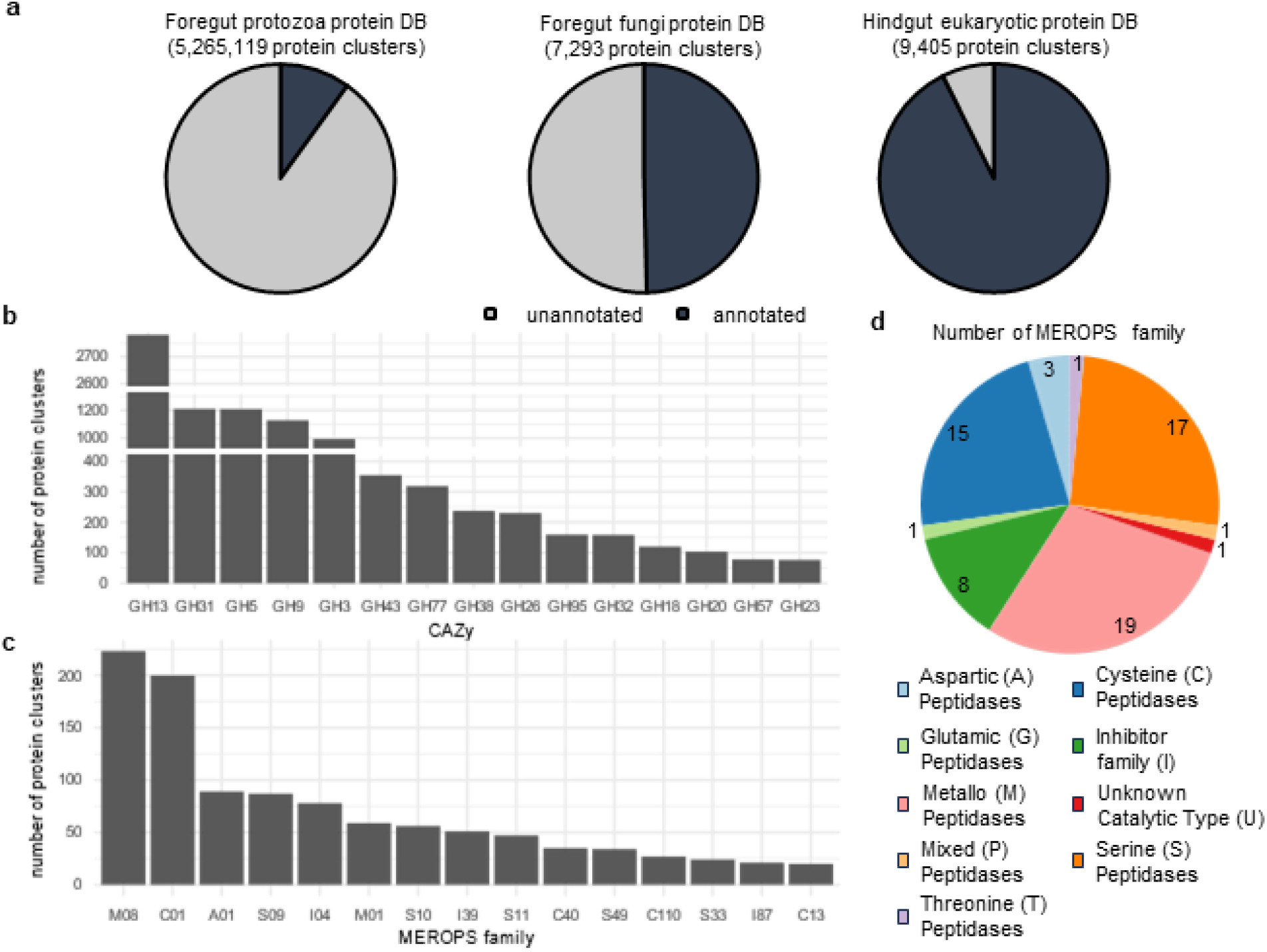
Diverse ruminant gut eukaryotic proteins identified in the databases. **a**, The number of protein clusters identified in foregut protozoal, foregut fungal, and hindgut eukaryotic protein databases. **b**, The most prevalent foregut protozoal glycoside hydrolases (GHs) identified. **c**, The most prevalent foregut protozoal secretory (with identified signal peptides) peptidases and peptidase inhibitors identified. **d**, the number of unique foregut protozoal secretory peptidases or peptidase inhibitors identified based on catalytic sites.

Previously, bacteria and fungi were credited for most of the cellulolytic activity within the rumen microbiome. However, a recent genomic study^17^ and a proteomic study^7^ have revealed a substantial proportion of genes encoding glycoside hydrolases (GHs), including cellulase and hemicellulases, within rumen protozoa^7^. In the present study, we annotated the foregut protozoal database for GHs. We found that amylases (e.g., GH13) and cellulases (e.g., GH5 and GH9) are among the most prevalent CAZymes identified (Fig. 5b), underscoring the significant fibrolytic and amylolytic capability of rumen protozoa.

Rumen protozoa are known for their ability to degrade protein, especially microbial protein. We thus also annotated the foregut protozoal database for peptidases. Among the peptidases identified, metallo, aspartic, cysteine, and serine peptidases were predominant, with M08, C01, A01, and S09 peptidases being among the most prevalent secretory peptidases (Fig. 5c). *Prevotella*, the most predominant genus of rumen bacteria with broad protease activity^29^, exhibits 48 different families of secretory peptidases^30^. In comparison, 58 different secretory peptidase families were identified from the rumen protozoal protein (Fig. 5d), highlighting their substantial proteolytic potential. Furthermore, the identification of ferredoxin hydrogenase of rumen protozoa (Supplementary Table 3) corroborates their ability to produce hydrogen.

### Ruminant gut eukaryotic protein databases improve protein identification in proteomics

Recently, a proteomic study demonstrated the utility of protozoal SAGs and genomes as a reference search database in annotating rumen protozoal proteins and assessing the significance of rumen eukaryotes in rumen functions^7^. Considering that reference search databases are a major bottleneck in metaproteomic studies of complex microbiomes and given the previously demonstrated greater genomic and functional diversity of the rumen eukaryotes, we evaluated the resource potential of expanding protozoal genomes with ruminant eukaryotic proteins identified with GutEuk to assist protein detection from rumen metaproteomic data. Using the same pipeline and threshold as the original aforementioned study^7^, we compared the number of identified protein groups from each sample with the original genome database (one *Entodinium caudatum* genome and 18 SAGs of rumen protozoa), the original genome database combined with the GutEuk-identified rumen eukaryotic search database resource (no additional SAGs included). The database expansion led to the identification of substantially more protozoal protein groups in both cow and goat rumen samples, resulting in a 64% increase in the number of protein groups identified in cow samples and a 20% increase in goat samples on average (Fig. 6). In some samples, around 200% increase was seen for the protozoa protein groups. This suggests that previously unknown protozoa are likely dominant in certain samples. Similarly, the expanded database greatly increased the number of fungal proteins identified, with a 118% and 70% increase in protein groups identified in cow and goat samples, respectively. However, on average, only 10 fungal protein groups were identified in the original study. It is also important to note that rumen samples were obtained via esophageal tubing for this particular metaproteomic study, meaning they were largely constituted of rumen fluid, which is known to have significantly lower fungal biomass than the solid fraction.

**Fig. 6:**
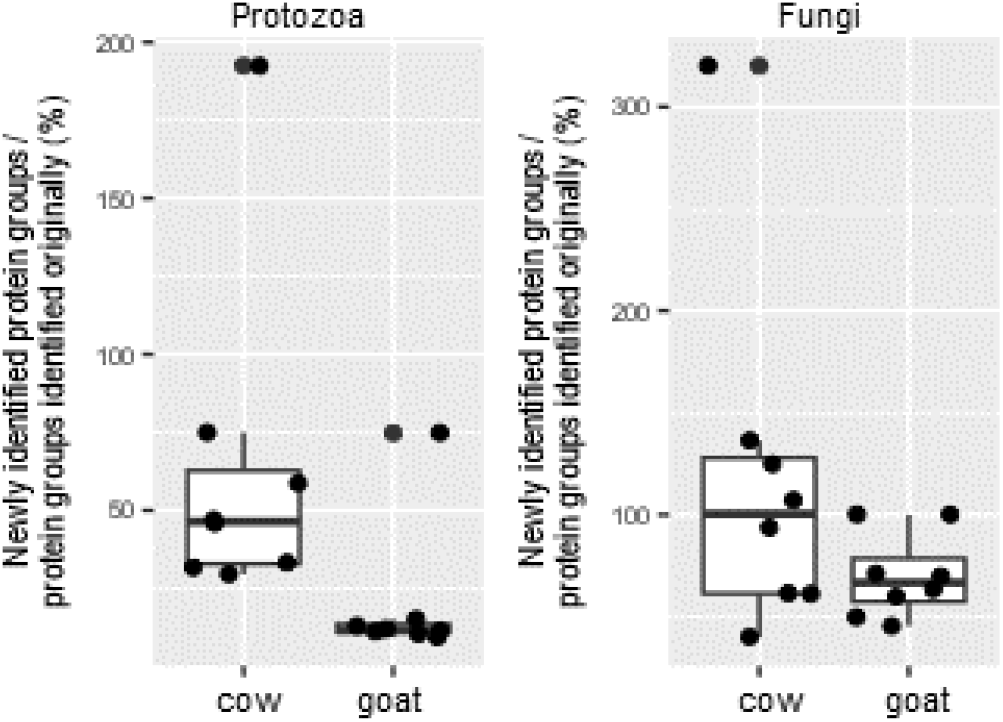
Increases in the number of rumen protozoal and fungal protein groups identified with the new protein databases compared with the previous rumen proteomic study. Box plots indicate the median (middle line), 25th and 75th percentile (box), as well as individual observations (dots).

## Discussion

Historically, gut eukaryotes have been studied primarily from a parasitological perspective. However, eukaryotic populations are also ubiquitous components of the gut microbiome^31,32^, and they contribute significantly to the overall microbial assembly, host nutrition, and physiology^33-35^. Their role is particularly significant in the rumen, the foregut of ruminants, where they can constitute up to 50% of the microbial biomass—a proportion much higher than in other ecosystems. These microbes play direct roles in rumen fermentation and fiber degradation^4^ and directly influence several important host characteristics, including feed efficiency and methane emissions^13^. Despite recent efforts in the cultivation and genomic sequencing of rumen protozoa and fungi, which is a highly labor-intensive process, we still lack a robust bioinformatic tool to analyze the eukaryotic microbes represented in shotgun metagenomes generated from whole rumen contents. Therefore, it is crucial to develop bioinformatic tools tailored to gut eukaryotic microbes, particularly those in the rumen, to obtain a more comprehensive representation of their genomes and proteomes. Such new tools will enable a shift from taxonomic profiling to functional profiling, allowing for a deeper understanding of gut eukaryotic microbes as well as whole gut microbiomes.

Tiara and EukRep have been utilized for eukaryotic sequence identification; however, they fail to accurately identify rumen eukaryotes. To facilitate metagenomic and functional analysis of gut eukaryotic microbes, especially those in the rumen, we developed GutEuk. This tool outperforms Tiara and EukRep in both precision and recall when identifying microbial eukaryotic genome fragments. The ensemble model implemented in GutEuk, which combines CNN and FNN, enables consistent performance for both short sequences (shorter than 5 kb) and long sequences (longer than 50 kb). The ability of GutEuk to further differentiate fungal and protozoal sequences is especially valuable for analyses of metagenomes derived from microbiomes that contain both types of eukaryotic microbes, such as the rumen microbiome. Although GutEuk was designed for metagenomes from the GI ecosystem, it also demonstrates consistent performance across genomes of diverse taxonomic lineages by precisely differentiating fungi and protozoa from other ecosystems. Since eukaryotic communities in the gut are less diverse than those in aquatic and terrestrial environments, and there is minimal overlap in the diversity of eukaryotic microbes among these ecosystems^31^, GutEuk is well suited for application in gut metagenomic studies.

Using GutEuk, we screened thousands of metagenomic bins assembled from the rumen and identified hundreds of individual eukaryotic bins. Our phylogenetic analysis of the dozens of identified protozoal bins with at least medium quality demonstrated the greater genomic diversity of rumen protozoa, particularly those in less-studied ruminant species. The relative sparsity of the recovered eukaryotic bins with high quality may be attributable to multiple factors, including the low abundance of eukaryotes in the rumen, their large and complex genomes, high genetic heterogeneity, and the presence of highly repetitive genomes^36^. Additionally, there is a risk of chimeric bins due to the presence of mobile elements and paralogous genes^21^. Annotation of eukaryotic bins should, therefore, be performed with caution. Binning of contigs into high-quality MAGs of eukaryotic microbes from metagenomic sequences, especially short sequencing reads, is probably challenging. By sequencing repetitive regions contiguously^36^, long-read sequencing, such as PacBio HiFi and Oxford Nanopore Technologies, may facilitate the assembling and binning of eukaryotic sequences from metagenomes, especially for protozoa that have highly fragmented small chromosomes, such as the *Entodinium* genus^28^. Additionally, ACR^37^, a bin refinement tool developed recently, may help facilitate genome-centric analyses of eukaryotes using metagenomics. Furthermore, similar to the assessment of prokaryotic genomes and MAGs, the completeness and contamination rates of eukaryotic bins should be determined based on the presence of lineage-specific marker genes and corresponding databases as implemented in BUSCO^38^ and EukCC^39^.

Several genomic structural features of eukaryotes, such as the exon-intron architecture and the lack of conserved splicing signatures, complicate the gene-calling process in eukaryotic genome annotations. To overcome these limitations, a reference- and homology-based pipeline, MetaEuk^40^, with high precision (>99.9%) was developed^40^. Utilizing this pipeline and a database combining single-cell protozoal transcriptomes and poly-A enriched rumen metatranscriptomes, we screened the contigs assembled from around one thousand ruminant GI metagenomes (see methods). This analysis led to the identification of millions of ruminant eukaryotic proteins, further highlighting the great metabolic potential of rumen protozoa, consistent with the previous studies^7,17^.

Leveraging the databases we compiled, we tested their resource applicability by reanalyzing the detected proteomic profiles of a recent study^7^. We substantially increased the number of identified protein groups from both fungi and protozoa. The additional proteins identified are primarily from the ruminant eukaryotic sequences identified by GutEuk, validating the genomic identification and protein prediction pipeline we used. The potential for chimeric binning is unlikely to confound the protein prediction as all the genomic sequences are predicted to be either fungal or protozoal. With the assistance of GutEuk and the derived protein database, we could better understand the distinct metabolic contributions of fungi and protozoa in the rumen, particularly under varying dietary and microbial interventions. However, protein prediction will require further genome curation, as described previously^21^, to attribute expressed protein to specific species.

Overall, the new tool, GutEuk, combined with the associated databases, enables comprehensive analyses of GI eukaryotes in metagenomes. The expansion of the reference genomes with proteins identified from rumen metagenomes enhances metaproteomic studies of the rumen microbiome. The expanded database also enhances read-based ecological analyses of gut eukaryotic microbes with tools such as EukDetect^41^, which provides taxonomic identification in samples with finer resolution compared to metataxonomics. GutEuk and the associated databases can help provide new insights into the functional contributions of ruminant fungi and protozoa to the rumen ecosystem as well as their interactions with diets and other microbes.

## Methods

### Evaluating existing tools in identifying and classifying rumen protozoa

Currently, Tiara^20^ and Eukrep^21^ are available tools for identifying eukaryotic contigs assembled from metagenomes. To evaluate their performance in identifying sequences of eukaryotes, we utilized the 52 rumen protozoa SAGs reported recently by Li, et al. ^17^. The telomere sequences were trimmed off from the telomere-capped contigs with a custom Python script. Since both tools were trained with contigs of 5kb length, the contigs of each SAG were chopped into 5kb fragments with a 1kb sliding window and used for benchmarking (see below).

### Genome dataset preparation

To specifically identify eukaryotic sequences in metagenomes derived from the gut, we compiled a dataset comprising genomes of prokaryotes, fungi, and protozoa. Briefly, 85,205 representative genomes of prokaryotes were downloaded from GTDB release 214.0 (https://data.gtdb.ecogenomic.org/releases/release214/214.0/). These genomes represent 80,789 species of bacteria and 4,416 species of archaea. Fungal genomes were downloaded from the MycoCosm genome portal (https://mycocosm.jgi.doe.gov/mycocosm/home). When multiple genomes were available for a fungal species, we only selected one genome with the highest quality. We also filtered out the genomes of mushrooms and soil fungi, reducing the number of fungal genomes not relevant to gut microbiomes. A total of 679 fungal genomes were retained, representing 639 genera. Protozoal genomes were downloaded from the NCBI GenBank database on September 2^nd^, 2023, and we selected the most complete genomes per species. We combined these protozoal genomes with the 52 rumen protozoal SAGs representing 19 species across 13 genera^17^. To remove potential sequence contamination from symbiotic prokaryotes, we retained only the contigs flanked by the identified telomere sequences of protozoa. Additionally, we selected only one species per genus from the fungal and prokaryotic genomes to prevent classification bias toward well-represented lineages. The dataset includes 20,739 prokaryotic genomes/contigs, 639 fungal genomes/contigs, and 380 protozoal genomes/contigs.

The genomes/contigs from each class were randomly assigned to one of the following datasets: training (70%), validation (15%), and testing (15%) to be used to train the classifier (see below). Within each dataset, genomes or contigs were randomly fragmented into 5 kb fragments with a sliding window of 5 kb. Then, we randomly selected 28.5% of the prokaryotic and fungal fragments and 75% of the protozoal fragments. This selection ensured a dataset size balanced with prokaryotic and gut eukaryotic sequences. The genomes/contigs used for training, validation, and testing are detailed in Supplementary Table 2.

### Development of GutEuk and training

We developed GutEuk, a two-stage classifier designed to differentiate prokaryotic and eukaryotic (fungal and protozoal) sequences in the first stage and then further differentiate fungal and protozoal sequences in the second stage. GutEuk employs an ensemble model comprising a feedforward neural network (FNN) and a convolutional neural network (CNN) in each stage (Fig. 1a) to classify individual input sequences after they were fragmented into 5kb length. For each input sequence, its 4-, 5-, and 6-mer frequencies were calculated, and its nucleotide bases (A, C, G, T) were one-hot encoded as [1, 0, 0, 0], [0, 1, 0, 0], [0, 0, 1, 0], and [0, 0, 0, 1], respectively). The *k*-mer frequencies and the one-hot encoded DNA sequences served as the input of FNN and CNN, respectively. In the first stage, the probability of a sequence being eukaryotic origin is calculated by FNN and CNN. The probabilities calculated by the two networks are then averaged. If the averaged probability of an input sequence exceeded 0.5, it was subjected to another round of FNN and CNN classification to be fungal or protozoal. Input sequences identified with a probability less than 0.5 were considered prokaryotic. For input sequences shorter than 5kb but longer than the user-defined minimal length, they were zero-padded to the right to reach 5kb, and their *k*-mer frequencies are calculated based on the original input sequences (Fig. 1b). For sequences longer than 5kb, the remainder fragments shorter than 5kb after the 5kb fragmentation were also zero-padded on the right and processed as described above. The origin of the input sequences (before fragmentation) was determined based on the predicted origin of their fragmented sequences. A user-defined confidence level was pre-set at the first and second stages (*pt*_*1*_ and *pt2*, respectively). If the percentage of fragments of the input sequence classified as either prokaryote or eukaryote was less than the threshold *pt*_*1*_, it would be classified as not available (NA). Otherwise, they were classified as either prokaryotic or eukaryotic based on the majority rule. Similarly, input sequences were further classified in the second stage if the percentage of their fragments classified as either fungal or protozoal exceeded the threshold *pt*_*2*_.

The CNN and FNN models in each stage were trained separately in PyTorch^42^ with the cross-entropy loss and AdamW optimizer. To optimize the hyperparameters, the model at different settings was manually curated and trained until the validation loss did not improve for ten consecutive epochs. The model with the best performance (in terms of validation loss) was used to predict sequence origins.

### Benchmarking EukRep, Tiara and GutEuk

To benchmark the performance of GutEuk against Tiara and EukRep, we filtered out the genomes originally used for training Tiara and EukRep in the testing set (2995, 93, and 43 genomes for prokaryotes, fungi, and protozoa, respectively, of diverse taxonomic origins). We first evaluated the performance of the three classifiers at the contig level. Because contigs vary substantially in length, the first 50 kb of those longer than 50 kb was chopped into fragments of 3, 4, 5, 8, 10, and 20 kb. Then, the performance of each tool was compared with the different lengths. For GutEuk, performance was also determined at different confidence levels. We further evaluated the performance of each tool at the genome level, with the criterion being the percentage of 5kb fragments of the genomes correctly classified.

### Identifying and functionally annotating microbial eukaryotic sequences from rumen metagenomes with GutEuk

We systematically analyzed 1,093 previously sequenced rumen metagenomes (Supplementary Table 1) to identify microbial eukaryotic sequences (Fig. 1c). Specifically, the raw reads downloaded from NCBI SRA were quality-filtered using fastp^43^ and assembled (individually for each metagenome) with Megahit^44^ (v.1.2.1) with the default setting. The quality-filtered reads were mapped to the resultant contigs with Bowtie2^45^, and the mapping results were used for binning with MetaBAT2^46^. Then, GutEuk was used to identify eukaryotic sequences from the resultant contigs. The bins greater than 3 MB were also screened with GutEuk with the parameter “-bin”. Based on the benchmarking result, we determined conservative thresholds to classify the contigs and bins. Specifically, in the first stage, bins were classified as either prokaryotes or eukaryotes only if 80% of the individual contigs within a bin were consistently classified as such and if the contigs were longer than 20 kb. In the second stage, eukaryotic bins were classified as fungi or protozoa only if more than 95% of the genomes were classified into the same category.

For functional annotations, the protein sequences from the previously sequenced polyadenylated-enriched rumen metatranscriptomics^17^ were first assembled using Plass^47^. The resultant protein sequences were clustered at 50% amino acid identity with MMseqs2^48^ to compile a rumen eukaryote protein sequence database. For the hindgut sequences, UniProtKB/Swiss-Prot release 2024_01^49^ was downloaded and used as the target database. Then, genes were predicted from the identified eukaryotic contigs and bins from the foregut and hindgut using MetaEuk^40^ with the clustered rumen eukaryotic protein sequences and UniProtKB/Swiss-Prot target database, respectively. The obtained protein sequences were functionally annotated using eggNOG-mapper v2^50^. Peptidases were also annotated from the identified protein sequences using DIAMOND^51^ with the e-value of 1e-10 as the threshold against the MEROPS database^52^. Signal peptides were identified from the protozoal peptidases using SignalP 6.0^53^, and those with a signal peptide were considered secretory.

### Phylogenomic analysis of recovered protozoa genomes

The completeness of the identified protozoa genomes/bins was assessed with BUSCO v5.6.1 by identifying lineage-specific single copy marker genes with the parameter “-l alveolata_odb10” applied. Only those with completeness exceeding 50% were subjected to further phylogenomic analysis. Briefly, the multiple sequence alignment of 30 BUSCO proteins of the identified protozoal genomes/bin and the 52 rumen SAGs^17^ were concatenated. Genomes/SAGs with less than 70% of these markers were excluded to ensure robust phylogenomic analysis. Specifically, the amino acid sequences of individual BUSCO proteins of the collected genomes were aligned with MAFFT v7.505^54^ and trimmed with trimAl v1.4^55^. The trimmed alignments of all the BUSCO proteins were then concatenated together with catfasta2phyml.pl (https://github.com/nylander/catfasta2phyml). The concatenated alignment was used to build a phylogenetic tree using IQTREE^56^ with the parameters “-redo -bb 1000 -m MFP -mrate E,I,G,I+G -mfreq FU -wbtl -nt AUTO” applied. The obtained tree was visualized with iTOL^57^.

### Evaluation of the new protein databases by reanalyzing the proteomics profiles from a recent rumen proteomic study

To benchmark the performance of the new protein databases in improving rumen eukaryotic protein identification and their applicability as a database resource for future analyses, we reanalyzed the metaproteomic data generated from both goat and cattle under two different dietary treatments using the same method described previously^7^. Briefly, FragPipe version 19^58^ was employed to analyze the raw data obtained from mass spectrometry (MS). The raw data were then analyzed with the original sample-specific databases used in the study or the original databases supplemented with the newly established protozoal (no additional SAGs included) and fungal protein databases (described in the previous section) using MSFragger^59^. To estimate false discovery rates (FDR), the databases were supplemented with contaminant protein entries such as human keratin, trypsin, and bovine serum albumin, alongside the reversed sequences of all protein entries. Variable modifications, including methionine oxidation and protein N-terminal acetylation, were accounted for, while carbamidomethylation of cysteine residues was held as a fixed modification. The choice of trypsin as the digestive enzyme allowed for a single missed cleavage, with matching tolerance levels set at 20 ppm for both MS and MS/MS. Filtering was executed to ensure a 1% FDR threshold. Quantitative analysis was conducted using IonQuant^60^. Only the proteins identified in the majority of the replicates in at least one of the treatments were considered present, as described in the original study.

## Data access

The established ruminant eukaryotic protein databases have been submitted to https://figshare.com/articles/dataset/ruminant_eukaryotes_protein_tar_gz/26210759. GutEuk, along with the codes used for visualization, are at https://github.com/yan1365/rumen_eukaryotes.

## Acknowledgments

This work is supported in part by the USDA National Institute of Food and Agriculture (Award number: 2021-67015-33393). We also thank the Ohio Supercomputer Center for providing the computing resources.

## Author contributions

M. Y. conceived this work, built the package, and performed bioinformatics analysis. T. A. performed the proteomics analysis. M. Y. and Z. Y. wrote the manuscript. All authors revised and approved the final manuscript.

